# Principles of cellular resource allocation revealed by condition-dependent proteome profiling

**DOI:** 10.1101/124370

**Authors:** Eyal Metzl-Raz, Moshe Kafri, Gilad Yaakov, Ilya Soifer, Yonat Gurvich, Naama Barkai

**Author notes:** These authors contributed equally to this work (Eyal Metzl-Raz, Moshe Kafri).

## Abstract

Growing cells coordinate protein translation with metabolic rates. Central to this coordination is ribosome production. Ribosomes drive cell growth, but translation of ribosomal proteins competes with production of other proteins. Theory shows that cell growth is maximized when all expressed ribosomes are constantly translating. To examine whether budding yeast function at this limit of full ribosomal usage, we profiled the proteomes of cells growing in different environments. We find that cells produce an excess of ribosomal proteins, amounting to a constant ≈8% of the proteome. Accordingly, ≈25% of ribosomal proteins expressed in rapidly growing cells do not contribute to translation. This fraction increases as growth rate decreases. These excess ribosomal proteins are employed during nutrient upshift or when forcing unneeded expression. We suggest that steadily growing cells prepare for conditions that demand increased translation by producing excess ribosomes, at the expense of lower steady-state growth rate.

## Introduction

Producing ribosomes is a major biosynthesis process in cells (Warner, 1999). Rapidly growing budding yeast, for example, contain about 190,000 ribosomes (von der Haar, 2008), each of which requires the production of over a hundred different proteins. Overall, ≈30% of the proteome of rapidly growing cells encodes for ribosomal proteins and ≈80% of all cellular RNA encodes for the rRNA components of the ribosomes. It is therefore expected that cells minimize the amount of ribosomes they produce, making only the ribosomes required for protein translation in the given condition.

The coordination of ribosomal protein expression with cell growth rate is of a particular interest. Ribosomes drive protein translation, and therefore cell growth. Increasing the amounts of ribosomes in cells could therefore increase growth rate. Yet, translation of ribosomal proteins competes with the production of other proteins, and could therefore introduce limitations in other essential cellular processes such as the production of metabolites needed for translation. Resource allocation models formalize this interplay, examining how to best distribute translation resources in order to maximize growth rate. These models predict that growth rate is maximized when all expressed ribosomal proteins are fully employed in protein translation (Bosdriesz et al., 2015; Dekel and Alon, 2005; Kafri et al., 2016a; Keren et al., 2013; Klumpp et al., 2013; Koch, 1988; Maaløe, 1979; Scott and Hwa, 2011; Scott et al., 2010, 2014; Vind et al., 1993). Indeed, producing excess ribosomal proteins that are not actively translating would compete with alternative cellular processes without contributing to protein production and cell growth.

The prevailing model therefore assumes that in growing cells, all ribosomes are constantly translating. However, recent studies suggest that while this model may hold in rapidly growing bacteria, slow-growing bacteria do contain excess ribosomes that are not actively translating (Dai et al., 2016). Further, we recently reported that in rapidly growing budding yeast, protein translation is not universally limiting for protein production and growth rate (Kafri et al., 2016b), raising the question of whether these cells employ all expressed ribosomes at full capacity, as predicted by resource allocation models.

Consistent with its major role in driving cell growth, the ribosome content of bacteria growing in different environments increases linearly with growth rate, independently of the specifics of each nutrient (Brauer et al., 2008; Bremer and Ehrenberg, 1995; Schaechter et al., 1958; Scott et al., 2010; Waldron and Lacroute, 1975; Warner, 1999; Zaslaver et al., 2009). This classical growth law was derived by measuring RNA content, which well approximates the amounts of ribosomes. In this work, we set out to extend these studies in three directions. First, we asked whether this bacterial growth law connecting ribosome content and growth rate is conserved in a model eukaryote, the budding yeast. Second, using proteomic profiling, we can now directly measure ribosomal protein levels and further compare their abundance with that of other protein groups. Finally, we extended the analysis also to non-steady state conditions, and in particular situations in which cells face increased translation demands, asking whether, and how, such conditions impact the relation between ribosome content and growth rate.

By analyzing the proteome composition of cells growing in a wide range of conditions, we show that the proteome fraction coding for ribosomal proteins scales linearly with growth rate, in a manner that is strikingly similar to that described in bacteria. Quantitative analysis of this scaling relation suggests that at each growth rate, a constant ≈ 8% fraction of the entire proteome encodes for an excess of ribosomes that are not actively translating. Accordingly, in rapidly growing cells, ≈25% of the ribosomal proteins produced are not employed in translation at a given time. This calculated inactive ribosome fraction significantly increases as the growth rate decreases, as verified in polysome profiling as well. We show that this excess of ribosomes is employed when cells are subjected to an unexpected increase in translation demands, for example upon a nutrient upshift or when forcing production of unneeded proteins. We suggest that steadily growing cells prepare for fluctuating conditions by producing excess ribosomes. This ribosomal excess enables a faster response to upshift conditions, but comes at the expense of limiting the cells’ instantaneous growth rate.

## Results

### The proteomic composition of budding yeast cells growing in different conditions

We employed mass-spectrometry analysis to define the proteomic composition of S. *cerevisiae* cells growing in three conditions: standard media (SC), media low in nitrogen and media low in phosphate. This data was combined with published proteomic datasets of budding yeast growing on 12 different carbon sources (Paulo et al., 2015, 2016). To avoid method-specific biases, all profiles were calibrated with an external data reference, defining absolute protein levels (Wang et al., 2012).

Examining the Pearson correlation between the different profiles (Figure 1A), we observed that the data is classified into two dominating clusters, depending on whether the carbon source provided supports fermentative or respiratory growth. Correlation between profiles within the same cluster was ≈0.7-0.9, while correlation between profiles assigned to different clusters was lower but still substantial (0.3-0.6). The data further demonstrated the induction of condition-specific proteins, as expected (e.g. activation of the phosphate and nitrogen starvation pathways), as well as differential expression of proteins involved in translation and stress response (Figure 1B).

**Figure 1:**
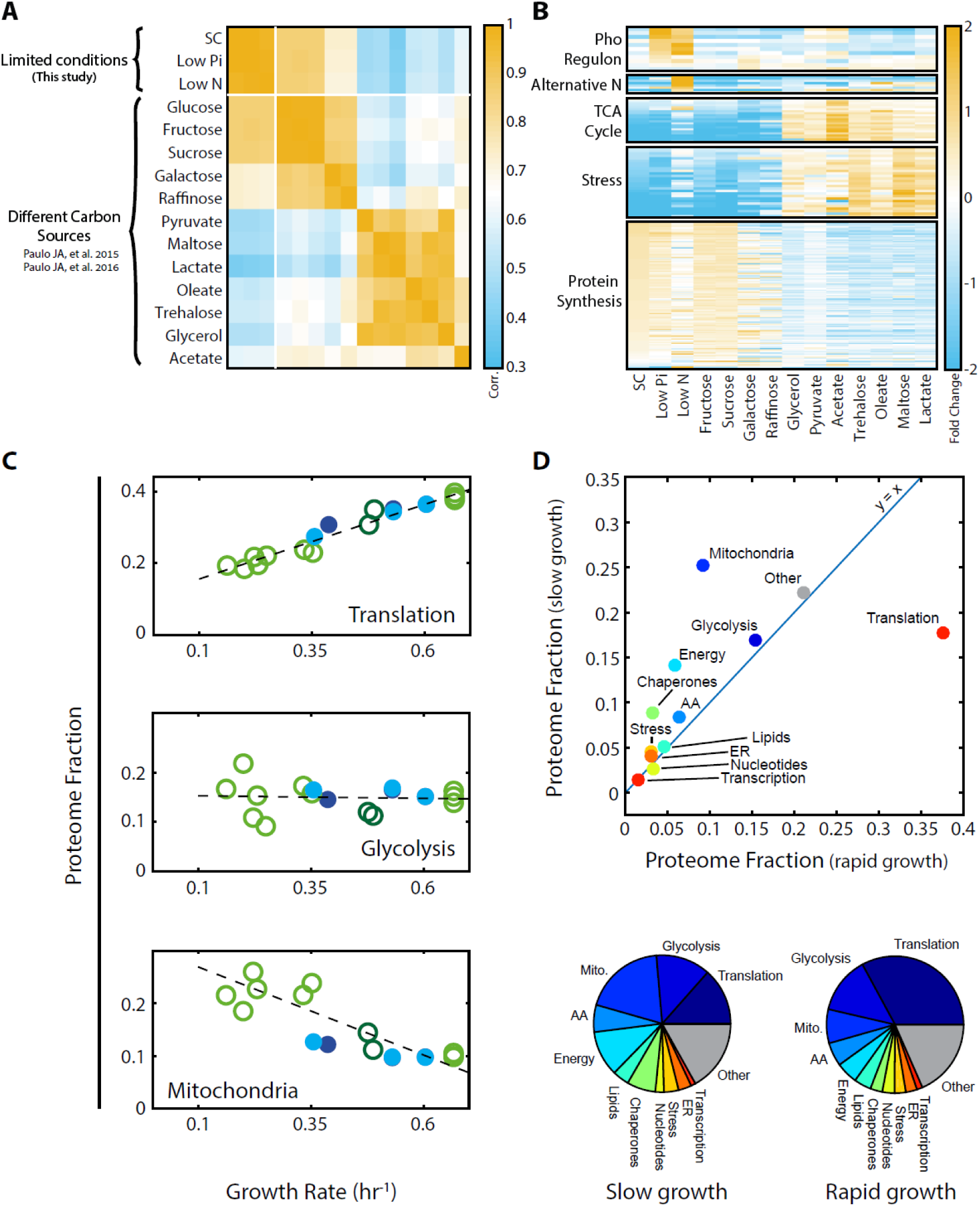
Proteomic analysis of budding yeast grown in different conditions. A. *Proteome profiles in our dataset clusters into two main groups:* Shown is the Pearson correlation matrix between proteome compositions in the indicated conditions. B. *Condition-dependent regulation of protein expression:* The expression of each protein in each condition was normalized by its mean expression over all conditions. Shown is the (Log2) protein expression of proteins in the indicated groups. See Supplementary Table 1 for protein names. C. *Expression of translation genes decreases in slow-growing cells:* Proteins were classified into eleven groups of related functions, which together included ≈80% of the proteome. For each condition, we calculated the fraction of the proteome coding for each of these eleven groups. Shown here are the proteome fractions of groups composed of proteins involved in translation, glycolysis or mitochondrial function, plotted as a function of cell growth rate. Additional protein groups are shown in Figure 1-figure supplement 1A. Filled circles correspond to data obtained in this work while empty circles are data from Paulo et al. (Paulo et al., 2015, 2016), as specified in Figure 2-figure supplement 1A. The proteins assigned to each protein group are specified in Supplementary Table 1. D. *The overall proteome composition in fast vs. slow growing cells:* The fraction of proteome encoding for each protein group was compared between the fast and slow growth condition in our dataset. In the upper panel, the proteome fraction encoding each specified protein group is plotted as a function of the proteome fraction encoding this group in the (averaged) slow growing condition. Fast growth (0.67 hr^−1^) corresponds to standard (SC) conditions, while slow growth (0.15 hr^−1^) is extrapolated from the linear fit to the abundance vs. growth rate relation, shown in (1C and Figure 1-figure supplement 1A). In the bottom panel, the same data is plotted as pie-charts.

Growth rate is a major determinant of proteome composition. Cells in the 15 conditions we examined grew at widely different rates, ranging from 0.15 to 0.65 generations per hour. To obtain a general overview of the changes in the proteome composition, we classified proteins into eleven groups of related functions, which together encompassed ≈80% of the proteome (Supplemental table 1), and examined how the relative abundance of proteins in each group changes with growth rate. In standard media, the proteome was dominated by translation-related factors (≈40%) and glycolytic proteins (≈15%) (Figure 1C). The fraction of glycolytic proteins remained largely invariant between conditions, while the translation-related fraction decreased with growth rate, reaching ≈15% in the slow-growing cells we examined. This decrease was accompanied by an increased abundance of condition-specific proteins, which in our dataset were mostly respiration-related. Mitochondrial proteins, for example, double in quantity to ≈20% of the proteome in the slow growing cells, reflecting the increased reliance of slow-growing cells on respiration (Figure 1C). The change in the proteomic fraction devoted to the different gene groups in slow versus fast growing conditions is highlighted by plotting the abundance of each group in the slowest growth condition, against its abundance in the fast growth conditions (Figure 1D, top) and by a pie-chart summarizing the proteome composition in those two conditions (Figure 1D, bottom).

### The proteome fraction encoding ribosomal proteins scales linearly with cell growth rate

To examine whether budding yeast follow the bacterial growth law connecting ribosome abundance with growth rate, we focused more specifically on the expression of ribosomal proteins. We defined the ribosomal fraction of the proteome by clustering together all proteins annotated as ribosomal subunits (Supplementary table 1). The proteome fraction coding for this group increased linearly with growth rate, irrespective of growth conditions, from ≈15% in the slowest growing cells to ≈30% in rapidly growing cells (Figure 2A). Therefore, budding yeast show the same growth law as was previously described in bacteria.

**Figure 2:**
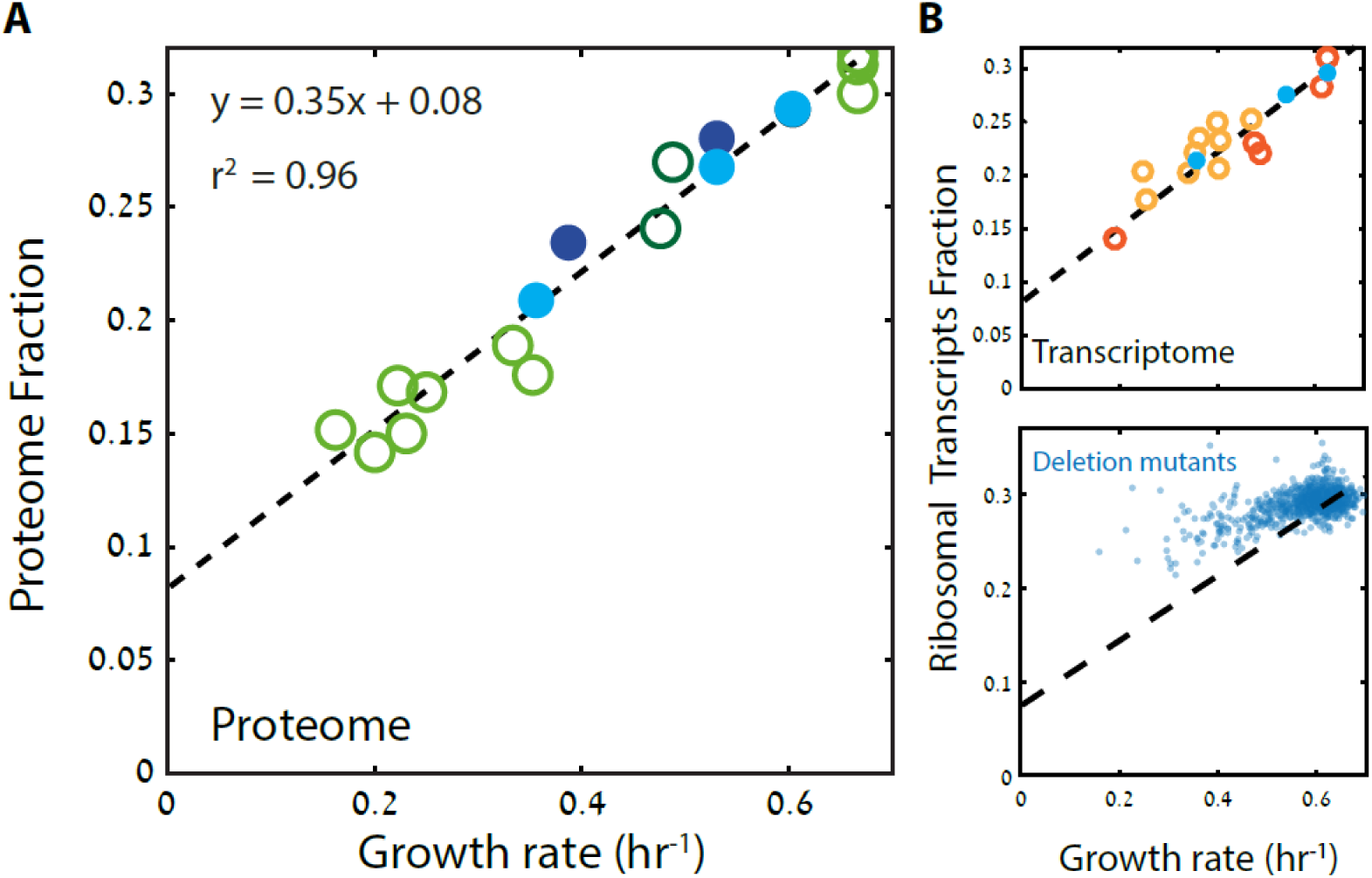
Ribosome content scales linearly with cell growth rate. A. *The proteome fraction coding for ribosomal protein scales linearly with growth rate:* Shown is the fraction of the proteome coding for the ribosomal proteins in each condition as a function of cell growth rate. Conditions are specified in Figure 2-figure supplemental 1A. Note the slope of this scaling curve Δμ/ Δr ≈ = 0.35x60min = 21mm^−1^. B. *The transcriptome fraction coding ribosomal proteins scales with growth rate:* Shown is the fraction of the ribosomal protein transcripts out of the full transcriptomes, as a function of cell growth rate. Conditions are specified in Figure 2-figure supplemental 1B. Dotted black line represents the linear fit in (A). C. *Mutant cells with reduced growth maintain high expression of ribosomal protein transcripts:* Same as (B) for data describing growth rate and transcription profiles of viable yeast deletion mutants (Kemmeren et al., 2014; O’Duibhir et al., 2014).

Cells may adjust their ribosome content by modifying protein translation, protein degradation or mRNA levels. To distinguish between these possibilities, we examined the transcription profiles of cells growing in the different conditions used for the proteome profiling. Notably, plotting the fraction of mRNA transcripts that code for ribosomal proteins as a function of cell growth rate showed the precise same quantitative scaling as observed at the level of the proteome (Figure 2B). Thus, regulation of the ribosomal protein fraction with growth rate occurs almost exclusively at the level mRNA transcription.

We asked whether the scaling between ribosome content and growth rate is strain-specific and regulated by changing growth conditions, or whether it also explains differences in growth rate between different strains growing in the same condition. We recently reported that budding yeast strains show a large strain variability in growth rate when growing on a pentose sugar (Tamari et al., 2014, 2016). Examining the transcription profiles of these strains, we find that the ribosomal fraction follows growth rate with the same qualitative scaling as observed when comparing growth rates in different conditions (Figure 2B, top). The general scaling relationship between ribosome content and growth rate is therefore conserved not only when comparing the same strain across conditions, but also when comparing differences between strains growing in the same condition.

We next asked whether the scaling of ribosome content and growth rate is also maintained in genetic perturbations that affect cell growth. To this end, we used a large dataset describing the gene expression profiles and growth rates of viable yeast deletion mutants (Kemmeren et al., 2014; O’Duibhir et al., 2014). Notably, in this case, the scaling between ribosome content and growth rate was largely lost (Figure 2B, bottom); in fact, most mutants that reduced growth rate had a minor effect on the expression of genes coding for ribosomal proteins. As a result, growth-affecting mutations expressed more ribosomal protein coding genes than expected given their growth rate. We observed a similar deviation from the scaling curve in cells forced to express a large amount of unstable mRNA (Figure 2-figure supplement 1D-E): here as well, the growth rate was largely decreased, with only a moderate decrease in the expression of ribosomal proteins. Therefore, while the scaling of ribosome content with growth rate is well conserved in wild-type strains growing in different conditions, it is largely absent in mutant cells. Many mutants actually maintained almost invariant ribosome content despite substantial reduction in growth rate, indicating that the scaling of ribosome content with growth rate result from cell signaling in response to the environment, rather than a direct causal relationship between ribosome availability and growth rate (Levy et al., 2007).

### The scaling of ribosome content with growth rate follows the predicted slope given by the ribosome translation time

To interpret the scaling relation between ribosome content and growth rate μ, we consider the general relationship between protein translation and growth rate characterizing exponential growth. Let us denote the fraction of the proteome encoding for ribosomal proteins that are active in translation by *r*_*a*_. In balanced exponential growth, the amount of cellular proteins precisely doubles at each cell division, captured by the specific growth rate μ Since proteins are translated by active ribosomes, this leads to the simple relationship μ = γ *r*_*a*_, where γ is the rate of ribosome translation (Kafri et al., 2016a; Maaløe, 1979; Scott et al., 2010).

Written differently, this relationship connects the two parameters we measure: growth rate μ and the proteome fraction encoding for translating and non-translating ribosomal proteins, *r:*

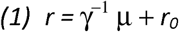

Here, *r*_*0*_ *= r-r*_*a*_, is the proteome fraction encoding for ribosomal proteins that are not actively translating. Note that this relationship Eq. 1, does not necessarily imply that ribosomal content scales linearly with growth rate, as in principle, the level of non-translating ribosomes *r*_*0*_ could change between conditions, leading to a non-linear relation between ribosome content *r* and the specific growth rate μ. The fact that linear scaling is observed, implies that *r*_*0*_ in fact remains constant, independent of growth conditions.

If the linear scaling we observe is indeed a reflection of balanced growth with a condition-independent *r*_*0*_ than the slope of the scaling curve, Δμ/Δ r should be given by γ, the rate of ribosome translation. We previously estimated that in budding yeast γ=23-24 min ^−1^ (Kafri et al., 2016b). Indeed, in our current measurements we find Δμ/Δ r ≈ 21 min^−1^, a value that is well in line with the predicted γ (Figure 2A-B). In bacteria, analogous measurements based on rRNA content predicted Δμ/Δ r = 6-10 min^−1^, consistent with the translation rate of the prokaryotic ribosome (Scott et al., 2010). Therefore, in yeast as well as in bacteria, ribosome content scales linearly with growth rate, with a non-zero intercept *r*_*0*_ and a slope whose value is given by the rate of ribosome translation.

### A constant fraction r_*0*_ ≈8% of the proteome encodes for non-translating ribosomal proteins

By comparing to the balanced growth rate we distinguish the fraction of ribosomal proteins that are actively translating *(r*_*a*_*=γr*_*a*_) from the inactive, non-translating part (*r*_*0*_). As we noted, *r*_*0*_ remains constant, independent of growth conditions. To better appreciate this, consider the zero-growth limit. In this limit, ribosomal proteins still account for *r*_*0*_ ≈8% of the proteome (Figure 2A-B). As protein translation in arrested cells is dramatically reduced, the majority of these ribosomes remain inactive. Indeed, arrested cells are known to maintain a pool of inactive ribosomes (Dai et al., 2016; van den Elzen et al., 2014).

The fraction of non-translating ribosomal proteins, *r*_*0*_, remains constant between conditions. Therefore, rapidly growing cells also devote *r*_*0*_ ≈8% of their entire proteome to producing ribosomal proteins not required for translation. This implies that at a given time, a substantial fraction of ribosomes are not actively translating: in rapidly growing cells ≈25% of the ribosomal proteins are inactive (*r*_*0*_≈8% relative to the total ribosomal fraction r≈30%), and this fraction increases with decreasing growth rate (Figure 3A).

**Figure 3:**
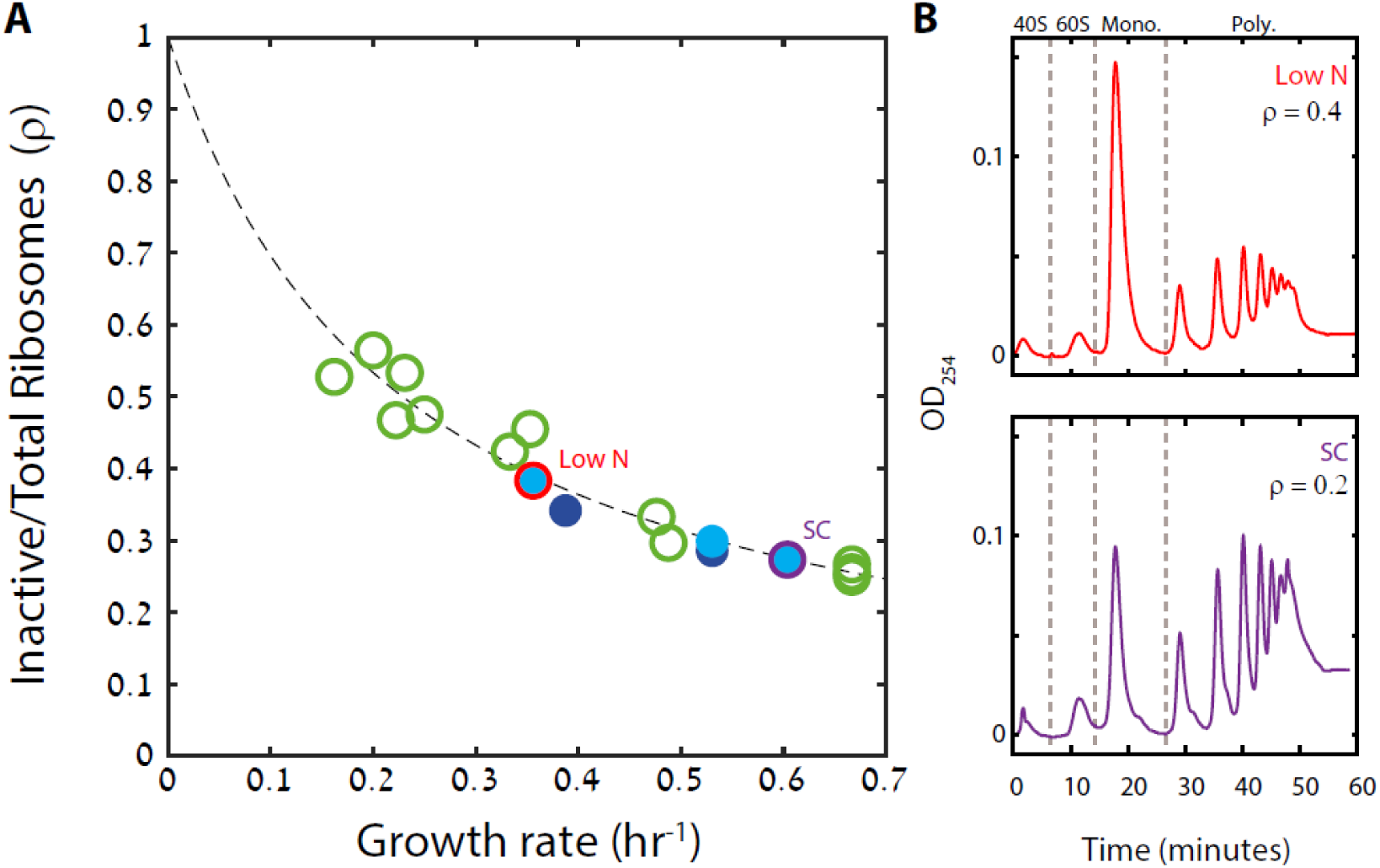
A substantial fraction of ribosomes is not actively translating at a given time. A. *The fraction on inactive ribosomes increase with decreasing growth rate:* Shown is the estimated fraction of inactive ribosomes (*ρ = r*_*0*_ / r) for each condition, as a function of cell growth rate. Conditions as specified in Figure 2-figure supplemental 1A. B. *Estimating the fraction of inactive ribosomes using polysomal profiling:* Cells were grown in the indicated conditions and their ribosomal content was analyzed on sucrose gradients as an indication for translational activity. Three independent experiments were performed, and representative profiles of raw data are shown. The fraction of inactive ribosomes was estimated by the ratio of monosomes (mRNAs bound by a single ribosome) to the total ribosome density and is indicated in the figures (*ρ*). Note the consistency of this value with the inactive fraction predicted by the proteome profiling analysis.

The fraction of inactive ribosomes is therefore predicted to be substantial even during rapid growth. As an independent approach to approximate this fraction, we used polysomal profiling, which quantifies the density of mRNAs bound by a different number of ribosomes. The most actively translating ribosomes are found in the polysome fractions (mRNAs bound simultaneously to multiple ribosomes). In contrast, monosomes are thought to contain primarily inactive ribosomes, although some of these may represent low-translating genes(Aspden et al., 2014; Heyer and Moore, 2016; Kelen et al., 2009). Examining the polysome profiles in cells growing in standard media (SC), low phosphate and low-nitrogen media, we observe a substantial increase in the monosome fraction of slow-growing cells provided with low nitrogen (Figure 3B). Differences in growth rate between SC and low phosphate were small and gave highly similar monosome density. Notably, the monosome fraction was also substantial in the fast growing cells (≈0.2). Further, in all the three growth conditions, the relative fraction of monosomes was largely consistent with the fraction of inactive ribosomes predicted from the scaling curve (Figure 3A-B)

### Cells employ the excess ribosomal proteins when preparing to enter stationary phase

Cells may use the excess ribosomes to accommodate an increase in translation demands. Entering stationary phase represents such a scenario (Ju and Warner, 1994), as the reduced availability of nutrients requires an increased production of glycolytic enzymes (Figure 1-figure supplement 1A, right column). To examine whether the fraction of non-translating ribosomes *r*_*0*_ decreases during this transition, we analyzed published proteomic profiles of batch-grown cultures entering stationary phase (Murphy et al., 2015). Indeed, ribosomal content decreases ≈2.5 generations (four hours) before growth rate drops (Figure 4A), resulting in a lower *r*_*0*_.

**Figure 4:**
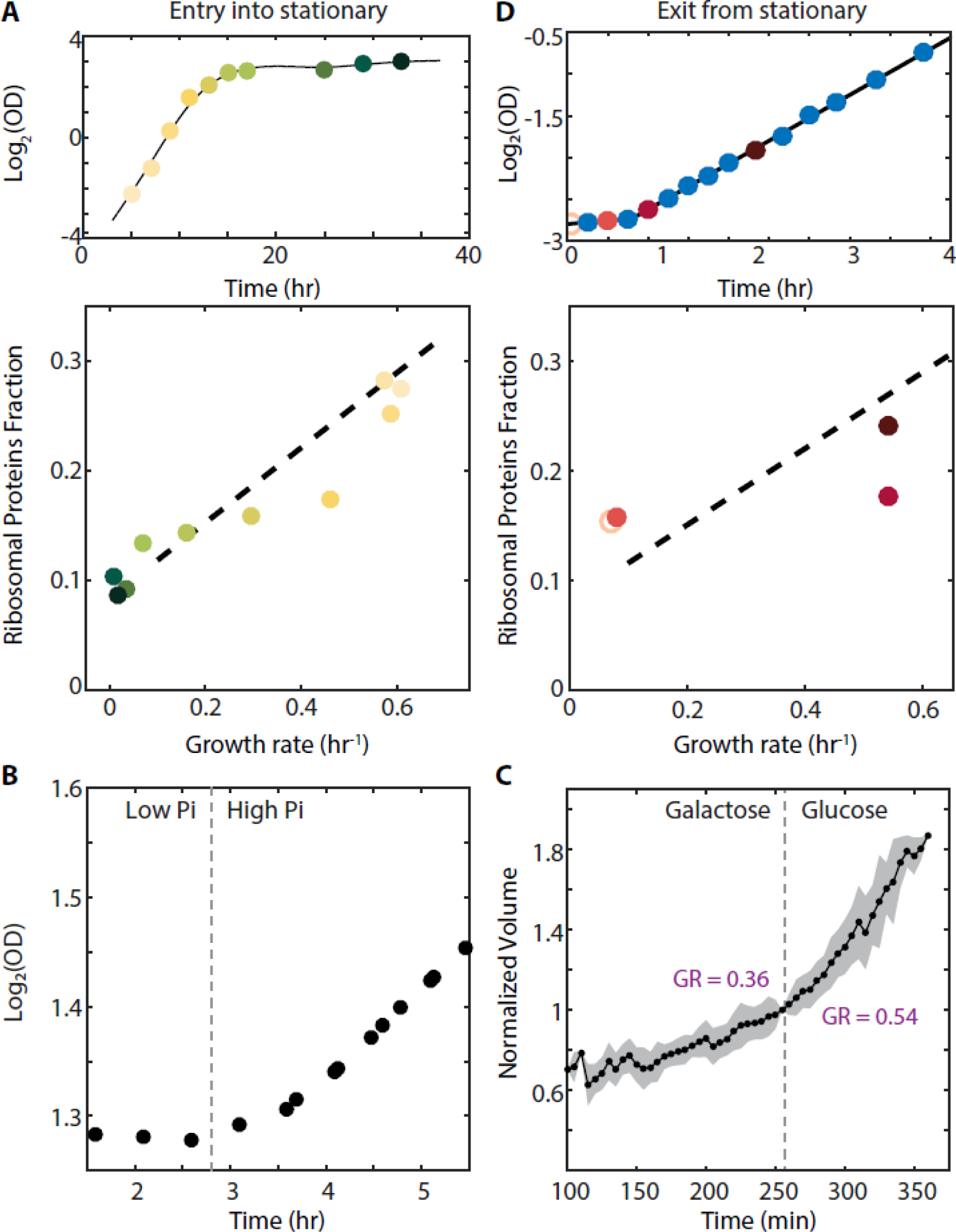
Cells employ a larger fraction of their ribosomes when subject to increased translation demands. A. *Cells entering stationary phase transiently decrease their predicted fraction of inactive ribosomes (r*_*0*_): Cells grown in batch culture were followed as they increase in density (top panel), and were subjected to proteome profiling at the indicated times. The bottom panel shows the fraction of proteome cording for ribosomal proteins along this time course. Data taken from Murphy et al. (Murphy et al., 2015). Color gradient from light to dark represents increasing time. B. *Cells growing in a phosphate-limited chemostat increase their growth rate immediately upon phosphate addition:* Continuous cultures were grown to steady state in a phosphate-limited chemostat at a dilution rate of 4.5 hr/gen. Media containing high phosphate was then injected into the growth chamber (dotted line). Shown is the density (OD) of the culture before and after phosphate injection. C. *Cells increase their volume growth immediately upon transfer to a preferred carbon source:* Cells were imaged using microfluidics-coupled live-cell microscopy, while their carbon source was changed from galactose to glucose (dotted line). Shown is the cell volume before and after the media change, averaged over all visualized cells. Specific growth rate was calculated 25 minutes before and after the upshift. D. *Cells exiting stationary phase resume rapid growth before increasing the ribosome content:* Cells were grown in SC until saturation (OD_600_ ≈6) and diluted back into fresh media at time 0 (empty circle). OD measurements were taken every 15-20 minutes as shown in the top panel, and samples for proteomic analysis were taken at the times marked in red. The ribosome fraction, as calculated from the proteomic data, is shown in the bottom panel as a function of cell growth rate. Each point is the median of three biological repeats.

Similarly, transcription profiling of cells growing in low phosphate showed an early decrease in ribosomal protein transcripts, which preceded the reduction in growth rate by about five hours (Figure 4-figure supplement 1A-B). Therefore, during preparation for stationary phase, cells reduce their overall ribosomal content but maintain a stable growth rate, suggesting that during this transient phase cells make a more efficient use of the available ribosomes.

### Cells employ the excess ribosomes during nutrient upshift

Translation demands increase when cells undergo nutrient upshifts. Under such conditions, nutrient limitation is lifted, and cells can resume fast growth practically immediately, provided that sufficient ribosomes are available to enable faster protein production. We examined the kinetics by which cells resume fast growth upon nutrient upshift by reanalyzing data we previously obtained. First, we found that continuous cell cultures growing in a phosphate-limited chemostat responded to a spike of high phosphate within minutes (Figure 4B). Second, using microfluidic-coupled microscopy, we followed individual cells transferred from galactose to glucose and observed that cells increase their specific growth rate from 0.36 hr^−1^ to 0.54 hr^−1^ within minutes of the transfer (Figure 4C). Therefore, cells increase their growth rate, and thus protein production rates, within a short time that appears insufficient for synthesizing new ribosomes, consistent with a more efficient use of available ribosomes and, more specifically, a decrease in the non-translating fraction *r*_*0*_.

To more directly correlate the ribosomal protein content and growth rate during nutrient upshift, we profiled the proteome of cells transferred from early stationary phase (OD_600_ ≈6) to fresh media (SC). Cells resumed rapid growth of 0.55 hr^−1^ following a ≈40 minute lag time. Proteome profiling revealed that ribosome content did not change during the lag time, but began to increase only after cells had attained fast growth (Figure 4D). Notably, during the initial growth phase, the excess of non-translating ribosomal proteins, *r*_*0*_, was dramatically reduced; In fact, to account for the observed growth rate during this initial growth phase, cells must have utilized the vast majority of ribosomes when initiating growth.

### The excess of non-translating ribosomal proteins is decreased when cells are forced to express unneeded proteins

We asked whether increasing translation demands during logarithmic growth will similarly decrease the fraction of ribosomal proteins that are not translating. To this end, we engineered cells to constitutively produce high mCherry protein amounts using a system we previously described (Kafri et al., 2016b). In this analysis, a library of strains expressing increasing amounts of mCherry proteins was generated, reaching ≈25% of the total proteome in the highest burden cells.

We asked whether this forced protein production would lead to a more efficient use of available ribosomes, reducing the proteome fraction *r*_*0*_ of the non-translating ribosomal proteins. Examining this requires measuring the scaling between ribosome content and growth rate in burden cells, as in Figure 2A. We therefore measured the growth rate and profiled the proteome for cells with different levels of mCherry burden in three conditions: standard media (SC), media containing low-phosphate and media containing low nitrogen. As expected, in all three conditions, growth rate and the ribosomal fraction *r* decreased with increasing *mCherry* levels (Figure 5A).

**Figure 5.**
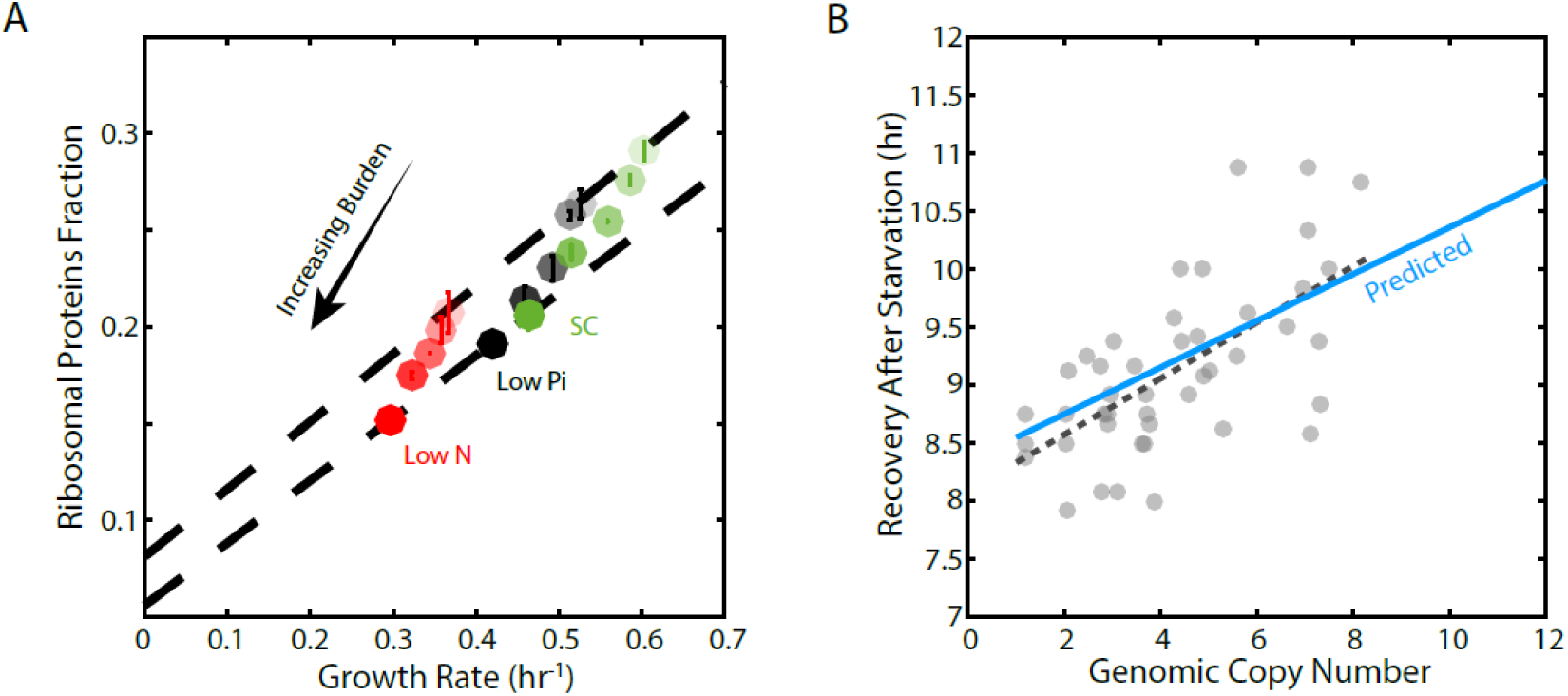
Forcing unneeded protein production reduces the pool of free ribosomes. A. *Scaling of ribosomal protein with growth rate in protein-burdened strains:* Five strains expressing increasing amounts of mCherry proteins were selected, and their proteome profiles and growth rates were measured in the three indicated conditions. Shown is the proteome fraction coding for ribosomal proteins in each strain and in each condition as a function of the cell growth rate. Different strains are indicated by the different shades of colors, with increased burden corresponding to a darker circle. Colors indicate the specific condition used. The two dashed lines correspond to the scaling curves defined by the no-burden and most highly-burdened strains: The top line is the same as in Figure 2A, while the bottom line describes the ribosome content of the highest burden as a function of its growth rate in the different conditions. B. *The reduced pool of inactive ribosome in the burden cells quantitatively accounts for their delayed exit from starvation-.* Cells expression different amounts of mCherry proteins were grown to saturation. The cells were kept in a weeklong starvation before dilution back to standard media. Recovery times (Y axis) were defined by the time at which cell density increased by 50%. Solid blue line represents the predicted recovery time based on the fold-reduction in free ribosomes, and dashed black line is the fit to the data. Data is re-plotted and analyzed from Kafri et al. (Kafri et al., 2016b).

For each level of burden, we plotted the ribosome content as a function of the cell’s growth rate in the three growth conditions examined (Figure 5A, Figure 5-figure supplement 1A). Similar to wild-type cells, in each of the burden strains, ribosome content scaled linearly with growth rate when compared across the three conditions. Notably, the slope of this scaling curve was independent of the level of the burden. Therefore, the ribosome translation rates γ (Figure 5A, dashed lines) remained invariant to the burden.

In contrast, the burden did affect *r*_*0*_, the predicted proteome fraction coding for non-translating ribosomes. This can be easily appreciated from the zero-growth limit. In this limit, ribosome content in high burden cells decreased from its wild-type value (*r*_*0*_=8.1%) to a lower value, *r*_*0*_*=*5.5% (Figure 5-figure supplement 1B). Therefore, the burdened cells employ a higher fraction of their ribosomes, leaving a smaller (but still condition-independent) pool of inactive ribosomes.

### The smaller pool of excess ribosomes in burden cells explains their delayed recovery from starvation

Burden cells exit starvation with a large delay (Kafri et al., 2016b; Shachrai et al., 2010). During this exit, cells must employ the excess ribosomal proteins *r*_*0*_ to initiate protein translation and cell growth. We therefore reasoned that the delayed recovery of the burdened cells may be due to the smaller fraction of residual ribosomes *r*_*0*_ they express.

To examine this prediction, we quantified the delay introduced by the burden when exiting starvation (Figure 5B; data from (Kafri et al., 2016b)). As expected, this delay increases linearly with the amount of mCherry. Notably, as predicted, the differences in recovery times were quantitatively explained by the difference in *r*_*0*_; for example, the recovery time of eight copies burden cells was prolonged by ≈15% compared to wild-type cells, consistent with the ≈15% decrease in the *r*_*0*_ in these cells.

### The fraction of ribosomes actively translating is tuned by growth conditions, but remains largely invariant to cell growth rate

A central implication of the scaling relation between ribosome content and growth rate (Figures 2A, 5A) is that when growth conditions change, cells tune not only the fraction of the proteome encoding for ribosomal proteins (*r*), but also the ratio between active and inactive ribosomes (*r*_*a*_ */r*_*0*_). As a direct consequence of the scaling law, this ratio increases linearly with growth rate when comparing wild-type cells grown in different conditions (Figure 6A). Comparing this ratio between wild-type and burden cells, however, we noted that in a given condition, the ratio was invariant to the burden. Thus, although burden cells grow more slowly, and express less ribosomal proteins compared to wild-type cells, the efficiency by which they employ the expressed proteins remains the same, depending only on the growth conditions. To verify this prediction, we used polysomal profiling. As predicted, the ratio of polysomes to monosome remains invariant between wild-type and burden cells growing in the same condition, despite their different growth rate (Figure 6B).

**Figure 6:**
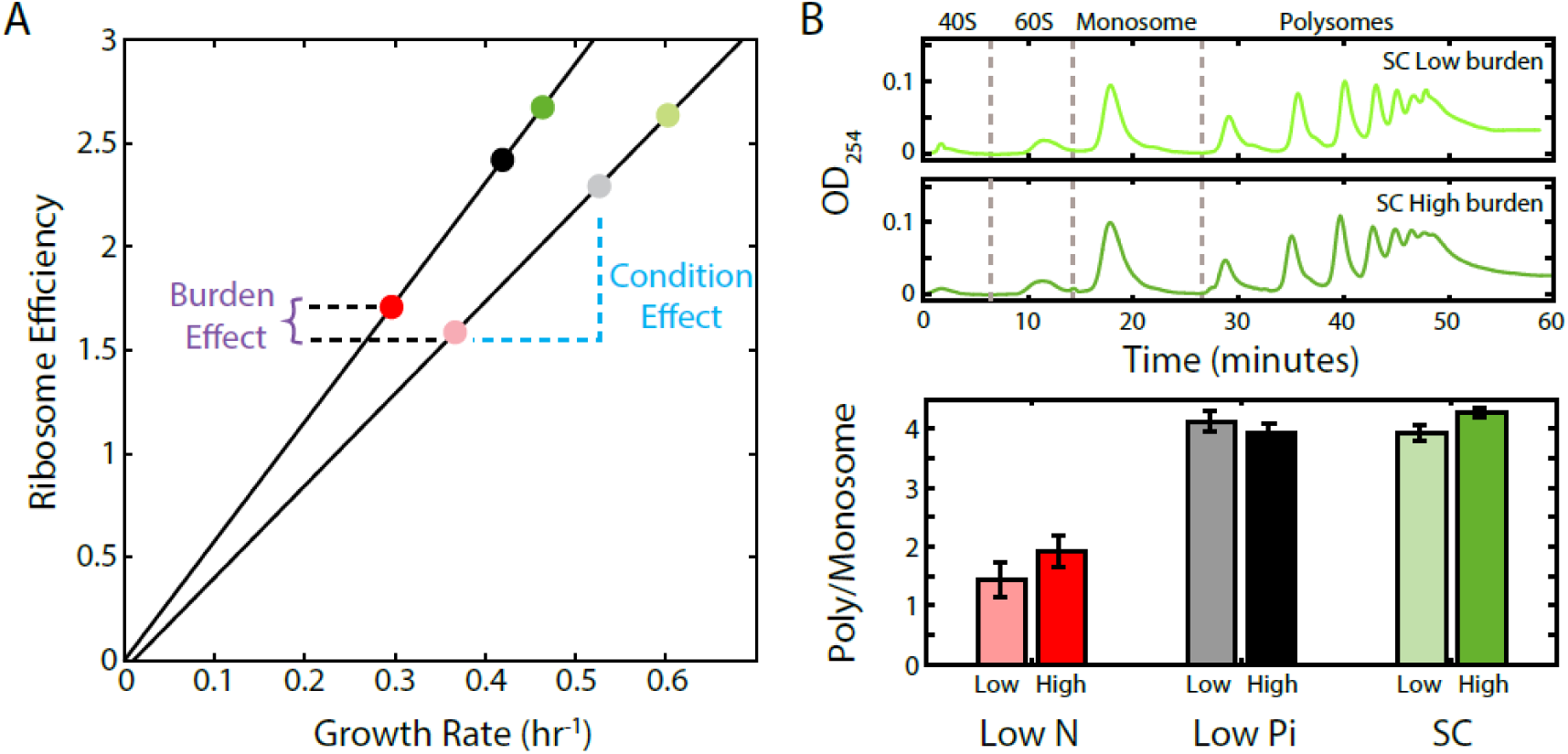
The ratio of active to inactive ribosomes remains invariant to protein burden. A. *The ratio of active to inactive ribosome predicted by proteomic data:* Shown is the ratio of active to inactive ribosomes *r*_*a*_*/r*_*0*_, as predicted from our analysis of proteomic data as a function of growth rate in wild-type and high-burden cells. Note that while this ratio decreases with growth rate when comparing the same stain across different conditions, it remains invariant to the burden when compared between the two stains within the same condition. B. *The ratio of active to inactive ribosome predicted by polysome profiling:* Low and high burden cells were grown in the indicated conditions and their ribosomal content was analyzed on sucrose gradients as an indication for translational activity. Representative profiles of raw data are shown on top. The ratio between the polysomes (active transcripts with more than one ribosome bound) to detached small and large subunits (40S & 60S) together with the monosomes (mRNAs bound by a single ribosome) is shown in a bar graph on bottom. SEM error bars are from three independent experiments.

Taken together with our results above, showing an invariant ribosome expression in slow growing mutants (Figure 2B, bottom), we propose that the tuning of ribosome content and of the fraction of ribosomes that are actively translating depend primarily on signaling from the environment with little contribution of growth-rate dependent feedback (Figure 7). Thus, although both parameters show a tight correlation with growth rate, this correlation is not direct, but rather results from evolutionarily tuned signaling.

**Figure 7.**
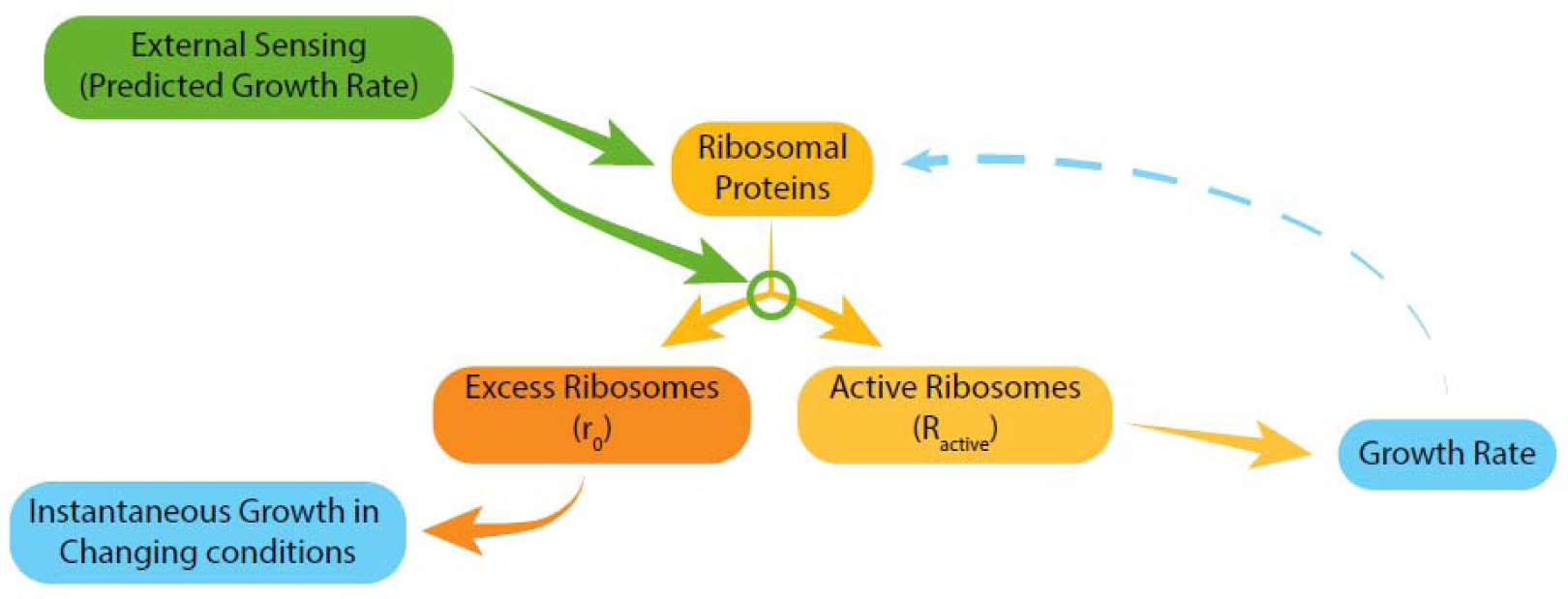
Model for ribosome allocation. Cells tune their ribosome content and ribosome efficiency based on signaling from the environment. Evolutionary tuning of this signaling results in a precise scaling of ribosome content with growth rate during logarithmic growth, but growth-rate dependent feedbacks play a minor role in the tuning of ribosome content or efficiency.

## Discussion

Ribosome production and function are the major resource-consuming processes in cells and its tight control in response to a variety of signals is well appreciated. Our study quantified a specific aspect of this control that is particularly relevant for understanding the overall regulation of cellular physiology: how the proteomic fraction coding for ribosomal proteins is regulated with growth rate. This question was extensively studied in theoretical models, and experimentally in bacteria, but received relatively little attention in eukaryotic models.

In bacteria, ribosome content is known to scale linearly with growth rate. Our proteomic analysis described an analogous growth law in budding yeast. Comparing steadily growing cells provided with different media, we find that the proteome fraction coding for ribosomal proteins scales linearly with growth rate. This scaling is strikingly similar to the growth-law described in bacteria: in both cases, the slope quantitatively matches the rate of ribosome translation and the scaling curve has a non-zero intercept.

Considering the fact that cells are growing exponentially, we have shown that the scaling curve we measured has one central implication - in any growth condition, a constant fraction (≈8%) of the proteome encodes for ribosomal proteins that are not actively translating at a given time. Accordingly, even rapidly growing cells still maintain ≈25% of their ribosomal proteins inactive. This result is surprising in light of the prevailing model, which assumes that cells tune their proteome composition in a way that ensures that all expressed ribosomal proteins are employed at full capacity. Such tuning is indeed intuitively expected: translation of ribosomal proteins competes with the production of proteins driving other cellular processes. Therefore, translation of ribosomes that do not contribute to translation, unnecessarily competes with other cellular processes. Why, then, would cells express extra ribosomes, more than needed for meeting translation demands?

We suggest that growing cells produce extra ribosomes in order to prepare for fluctuating conditions. Consider a nutrient upshift during growth in poor media: when the nutrient limitation is lifted, cells can increase their growth rate, provided that sufficient ribosomes are available. If cells were to precisely tune their ribosomal content to express only the ribosomes needed for the slow growing conditions, their level would limit the speed in which rapid growth could be resumed upon nutrients supply. Indeed, we have shown that cells employ the extra pool of expressed ribosomes when subjected to nutrient upshift, and also when subjected to other conditions inflicting an unexpected increase in translation demands. In particular, forcing cells to express high levels of unneeded proteins reduces the fraction of the inactive ribosomes during steady state growth, and, perhaps as a consequence, proportionally delays the exit of these cells from starvation.

Previous studies described strategies by which cells prepare for changing conditions, including the activation of stress genes by moderate stresses (Gasch et al., 2000; Guan et al., 2012; Levy et al., 2011; Mitchell et al., 2009) and the transition of a subpopulation into persistence (Balaban et al., 2004; Levy et al., 2012; Soll and Kraft, 1988; Yaakov et al., 2017).

Our study shows that this requirement to be prepared for fluctuating conditions affects not only specific genes or pathways, but also the central cellular processes such as the coordination of ribosome production with growth rate.

## Acknowledgments

We thank J. Paulo for sending unpublished growth data, G. Jona for assistance with the chemostats and our lab members for fruitful discussions. This work was supported by the ERC and the ISF.

---

**Figure 1-figure supplement 1:**
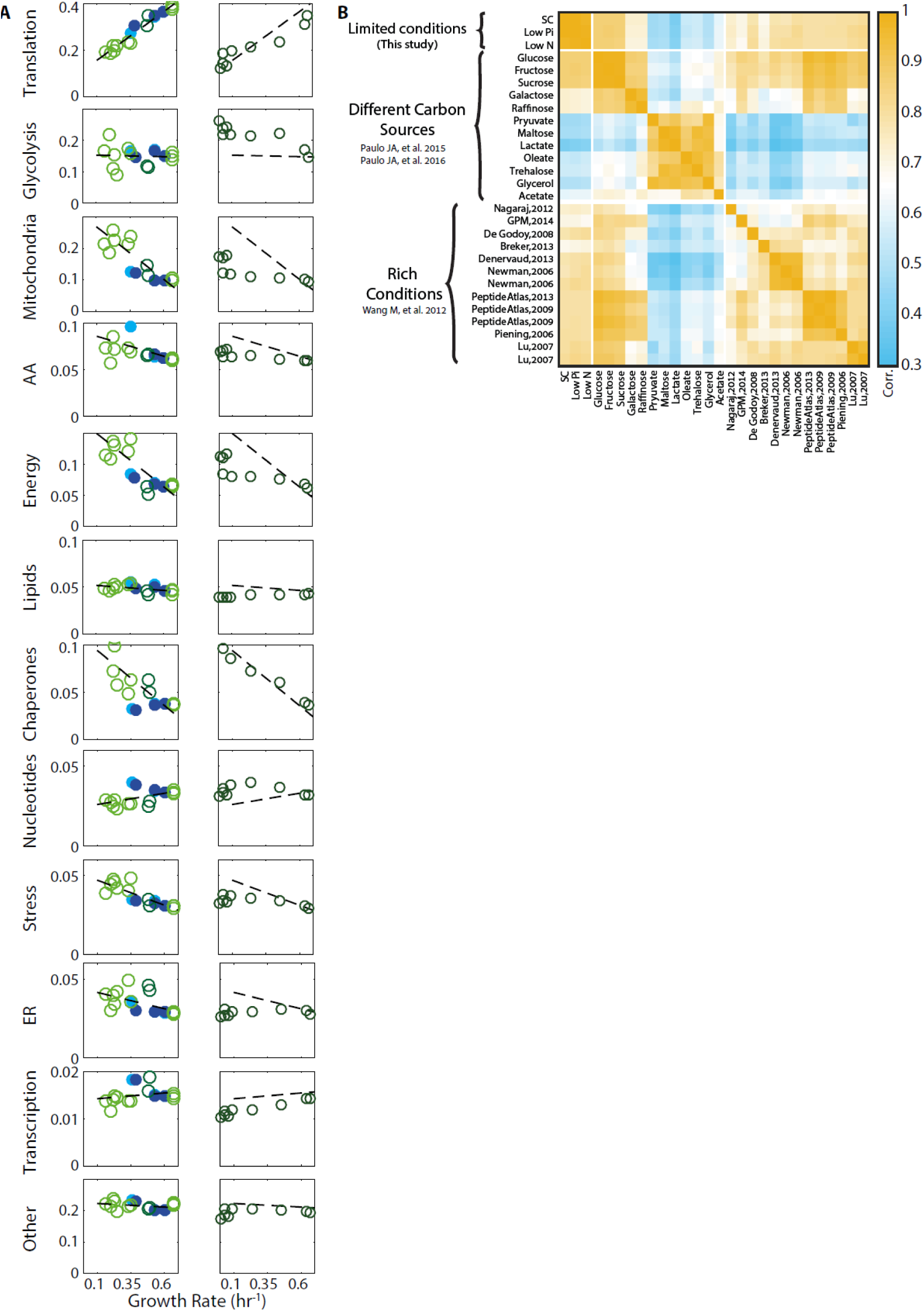
Proteome composition. A. The proteome composition under different conditions: Left column - proteome composition of all protein groups analyzed as indicated in Figure 1C. Right column - identical protein groups, with each circle representing a time-point along the growth curve as described in Figure 4A. B. Correlation between proteome profiles of cells growing in different conditions: Pearson correlation matrix as in figure 1A, including all rich condition proteomic datasets used in this study.

**Figure 2-figure supplement 1:**
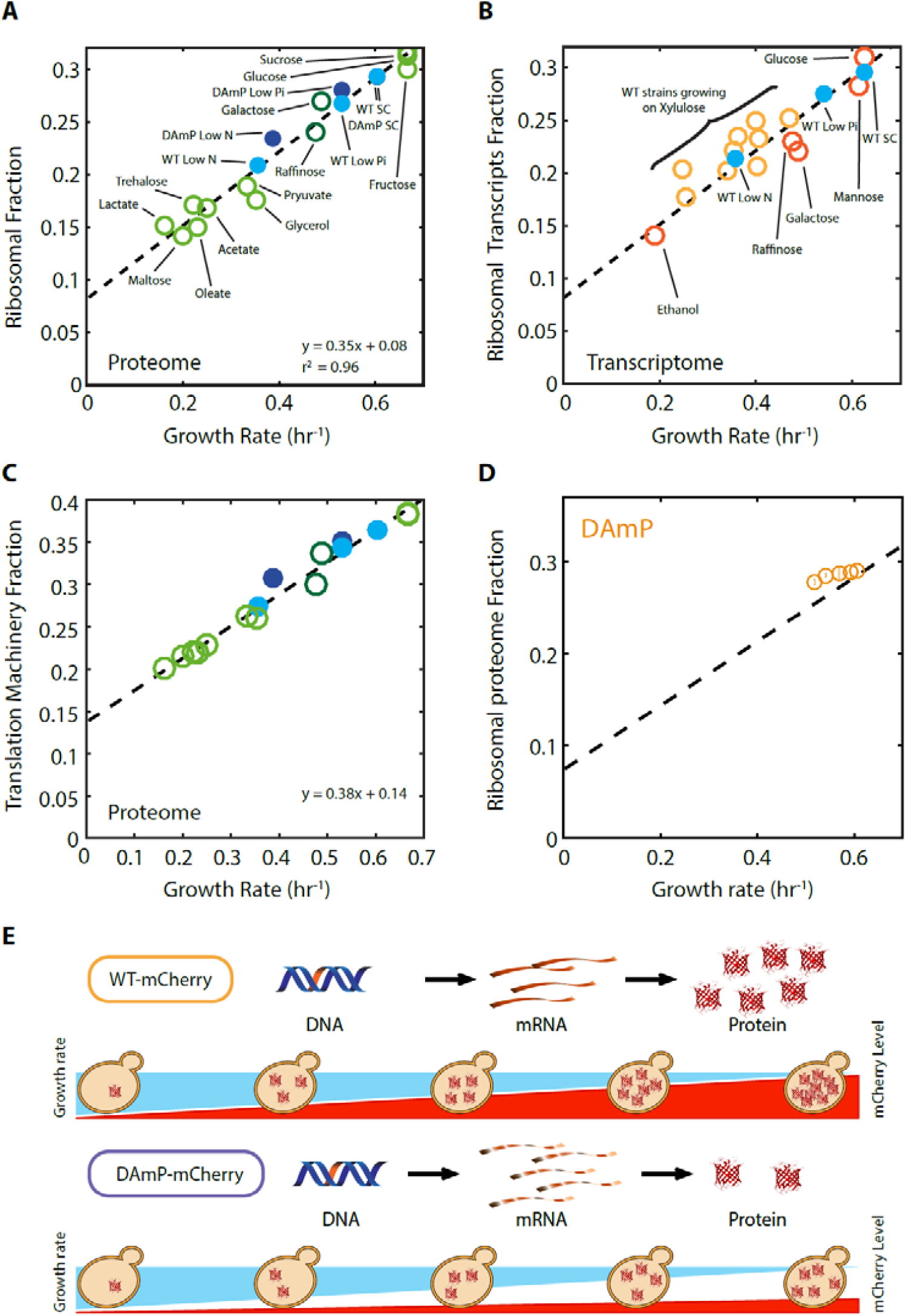
The proteome profiles of budding yeast cells growing in different conditions. A. Ribosomal fraction as a function of cell growth rate: Same as figure 2A, specifying the condition. B. Same as figure 2B, specifying the condition. C. Same as (A) for a broader definition of ribosome-associated proteins, see supplemental table S1 for full list. D. Ribosomal proteome fraction from cells forced to transcribe unstable DAmP transcripts are shown (Kafri et al., 2016b). Error bars represent the standard deviation around the median between 3 repeats. E. Generation of libraries of burdened cells. See (Kafri et al., 2016b).

**Figure 4-figure supplement 1:**
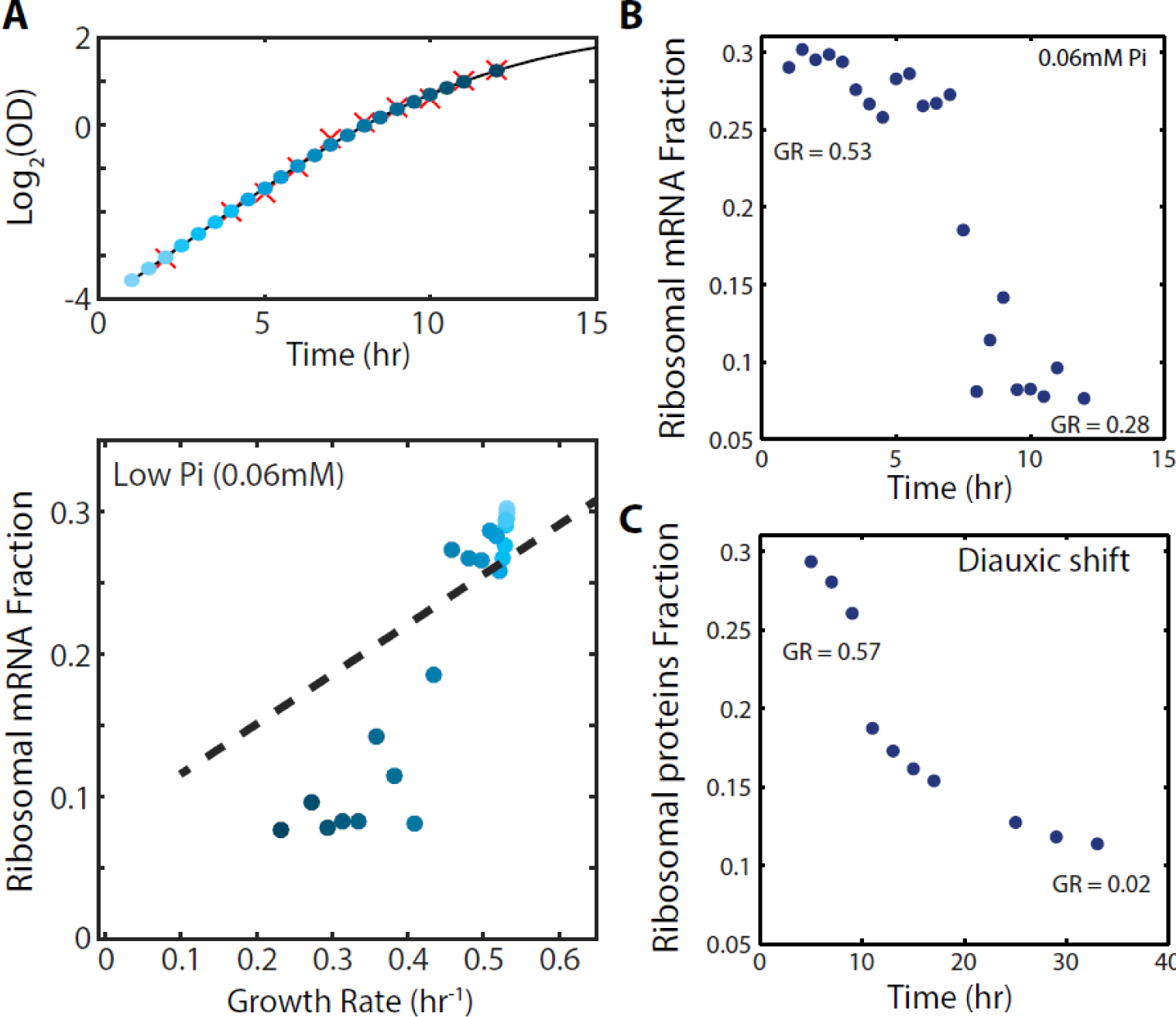
Cells can employ the excess ribosomes. A. The fraction of the transcriptome encoding ribosomal proteins in cells growing in low phosphate: Exponentially growing cells in SC media were diluted into low phosphate media (0.06mM). Samples were taken for OD measurements only (red marks, top panel) or OD measurements together with RNA samples (blue circles, top panel). The bottom panel shows the ribosomal fraction as calculated from the transcriptomic data from samples of the corresponding color in growth curve. B. Ribosomal expression data as in (A) plotted as a function of time. C. Proteomic ribosomal data as in Figure 4A plotted as a function of time.

**Figure 5-figure supplement 1:**
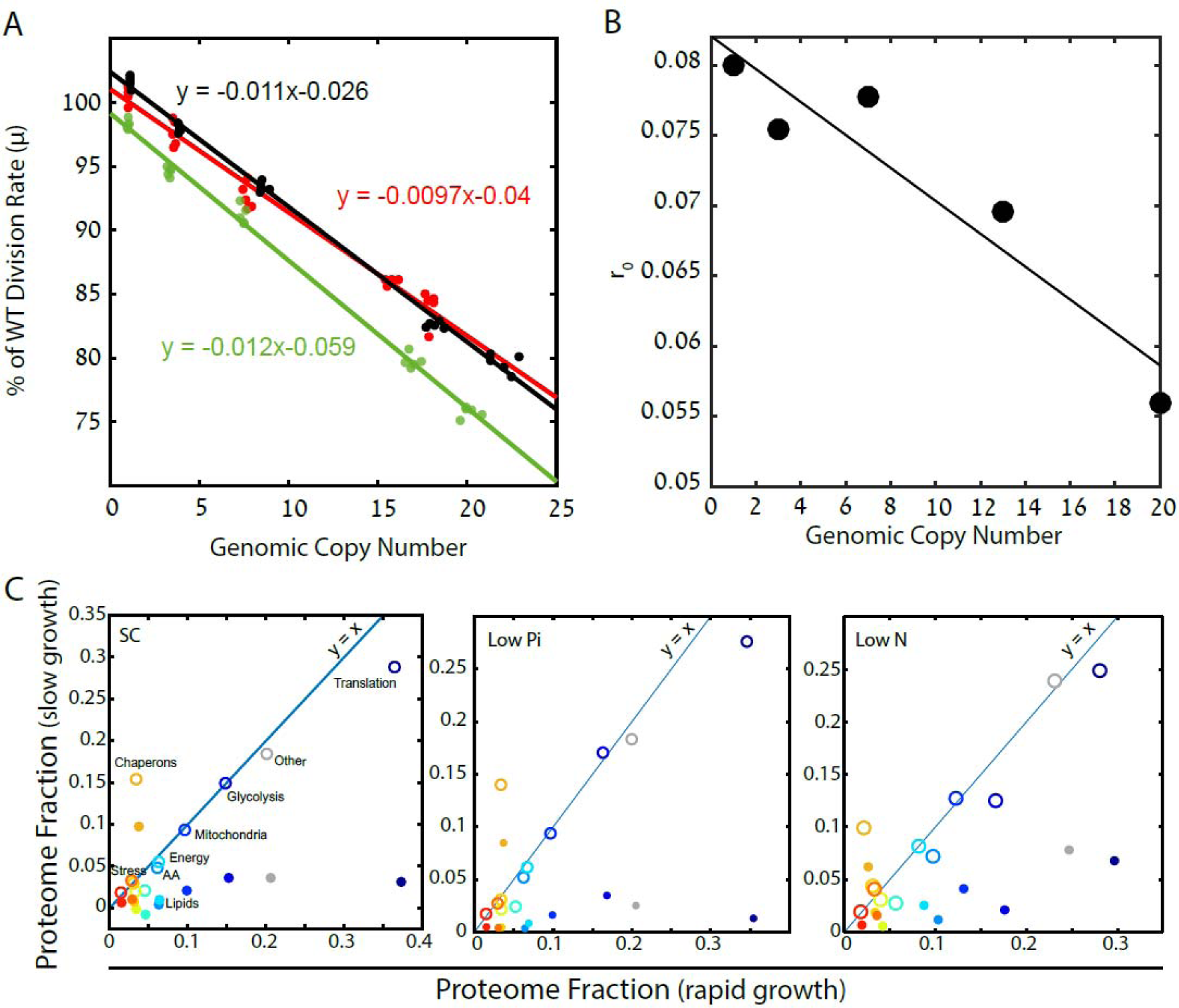
Cells forced to produce unneeded proteins have a smaller pool of excess ribosomes. A. Relative growth rates of burdened cells: The growth effect of mCherry copy number in different conditions is plotted; black low pi, green SC; red low N. Relative growth rate was calculated using competition experiments. See also (Kafri et al., 2016b). B. Excess ribosomes (r_0_) decrease with burden: The r_0_ fraction is plotted as a function of mCherry copy number, with a linear fit in black. This fit was used for calculations that took into account r_0_ of burdened cells. C. Burden passively impacts the ribosomal proteomic fraction: The fraction of proteome in burdened cells dedicated to the indicated protein group in fast (one mCherry copy) or slow (high burden) growing cells. Slow growth is extrapolated from the linear fit of increasing copy number. Closed circles represent all proteomic data including mCherry amounts; Open circles show only endogenous protein content.

## Supplementary Materials

### Materials and Methods

#### Media and strains

All budding yeast (S. *cerevisiae)* strains in this study were based on BY4742 (Brachmann et al., 1998) or Y8205 (Tong and Boone, 2007) laboratory strains. TDH3p-mCherry (protein fusion) was made by fusing the mCherry cassette to the end TDH3 protein using a standard PEG;LiAC;ssDNA transformation protocol (Gietz and Woods, 2002). The rest of the strains were constructed previously (Kafri et al., 2016b).

Strains were grown in SC medium or in SC medium depleted of a specific nutrient, as described in the main text. SC limiting media was prepared from YNB without the relevant nutrient (Low Phosphate medium - ForMedium, CYN0804, Low Nitrogen medium - BD 3101130). Phosphate depleted medium was prepared by adding phosphate in the form of KH_2_P0_4_ to a final concentration of 0.2mM. The level of potassium was preserved by adding KCL (instead of KH_2_P0_4_). Nitrogen limiting medium was prepared from YNB without amino acids and ammonium sulfate (BD 3101130) by adding separately 50 μM of ammonium sulfate and the essential amino acids.

#### Growth rates

Growth rates were calculated from the data reported previously (Kafri et al., 2016b). We further validated the WT growth rates in the various conditions by log-phase competition assays, as described below.

#### Log-phase Competition

Cells were grown overnight to stationary phase in the relevant media. GFP and mCherry strains were then co-incubated in the specified media at 30°C. The initial OD was set to ≈0.05, and the WT initial frequency was ≈50% of the total population. The number of generations was calculated from the dilution factor. Frequencies of GFP versus mCherry cells were measured by flow cytometry. The cells were diluted every ≈8 hours. Experiments were done with WT strains expressing GFP vs. mCherry burdened cells. A linear fit of the log_2_ for the WT frequency was used to calculate the slope for each competition. The relative fitness advantage is calculated from the slope divided by log_2_. The slope of the strains’ fitness advantage and their copy number were used to calculate the growth effect per burden copy.

#### Protein measurements

##### Sample preparation

All chemicals are from Sigma Aldrich, unless stated otherwise. Samples were subjected to in-solution tryptic digestion using a modified Filter Aided Sample Preparation protocol (FASP). Sodium dodecyl sulfate buffer (SDT) included: 4%(w/v) SDS, 100mM Tris/HCL pH 7.6, 0.1M DTT. Urea buffer (UB): 8 M urea (Sigma, U5128) in 0.1 M Tris/HCL pH 8.0 and 50mM Ammonium Bicarbonate. Cells were dissolved in 100 μL SDT buffer and lysed for 3min at 95°C and then centrifuged at 16,000 RCF for 10min. 100ug total protein were mixed with 200μL UB and loaded onto 30 kDa molecular weight cutoff filters and centrifuged. 200 μl of UB were added to the filter unit and centrifuged at 14,000 x g for 40 min. Alkylation using 100μl IAA, 2 washed with Ammonium Bicarbonate. Trypsin was then added and samples incubated at 37°C overnight. Additional amount of trypsin was added and incubated for 4 hours at 37°C. Digested proteins were then spun down to a clean collecting tube, 50ul Nacl 0.5M was added and spun down, acidified with trifloroacetic acid, desalted using HBL Oasis, speed vac to dry and stored in -80°C until analysis.

##### Liquid chromatography

ULC/MS grade solvents were used for all chromatographic steps. Each sample was loaded using split-less nano-Ultra Performance Liquid Chromatography (10 kpsi nanoAcquity; Waters, Milford, MA, USA). The mobile phase was: A) H20 + 0.1% formic acid and B) acetonitrile + 0.1% formic acid. Desalting of the samples was performed online using a reversed-phase C18 trapping column (180 μm internal diameter, 20 mm length, 5 μm particle size; Waters). The peptides were then separated using a T3 HSS nano-column (75 μm internal diameter, 250 mm length, 1.8 μm particle size; Waters) at 0.35 μL/min. Peptides were eluted from the column into the mass spectrometer using the following gradient: 4% to 22%B in 145 min, 22% to 90%B in 20 min, maintained at 95% for 5 min and then back to initial conditions.

##### Mass Spectrometry

The nanoUPLC was coupled online through a nanoESI emitter (10 μm tip; New Objective; Woburn, MA, USA) to a quadrupole orbitrap mass spectrometer (Q Exactive Plus, Thermo Fisher Scientific, Bremen, Germany) using a Flexion nanospray apparatus (Proxeon). Data was acquired in DDA mode, using a Top20 method. MS1 resolution was set to 70,000 (at 400m/z) and maximum injection time was set to 20msec. MS2 resolution was set to 17,500 and maximum injection time of 60msec.

##### Data processing and basic analysis

Raw data was imported into the Expressionist^®^ software (Genedata) and processed as described previously (Shalit et al., 2015). The software was used for retention time alignment and peak detection of precursor peptides. A master peak list was generated from all MS/MS events and sent for database searching using Mascot v2.5 (Matrix Sciences) and MSGF+ (Integrative Omics, https://omics.pnl.gov/software/ms-gf). Data was searched against the Saccharomyces cerevisiae (strain ATCC 204508 / S288c) protein database as downloaded from UniprotKB (http://www.uniprot.org/), and appended with 125 common laboratory contaminant proteins as well as the mCherry protein sequence (Uniprot accession X5DSL3). Fixed modification was set to carbamidomethylation of cysteines and variable modifications were set to oxidation of methionines and deamidation of N or Q. Search results were then filtered using the PeptideProphet algorithm (Keller et al., 2002) to achieve maximum false discovery rate of 1% at the protein level. Peptide identifications were imported back to Expressionist to annotate identified peaks. Quantification of proteins from the peptide data was performed using an in-house script (Keller et al., 2002). Data were normalized based on the total ion current. Protein abundance was obtained by summing the three most intense, unique peptides per protein. A Student’s t-Test, after logarithmic transformation, was used to identify significant differences across the biological replica. Fold changes were calculated based on the ratio of arithmetic means of the case versus control samples.

##### Proteomic datasets

We analyzed the following external mass-spectrometry datasets: two carbon sources (Paulo et al., 2015, 2016), diauxic shift (Murphy et al., 2015) and arsenate data sets (Guerra-Moreno et al., 2015). In each dataset, we normalized our glucose condition (SC) or time zero to the literature (median of 13 rich conditions from PaxBD (Wang et al., 2012), see conditions in Figure S1B, ‘Rich Conditions’). For the analysis, we used only the genes also present in our dataset. The carbon source data set (Paulo et al., 2015, 2016) growth rates were acquired from the papers or via personal communication. The diauxic shift data set (Murphy et al., 2015) growth rates were calculated from Figure 4A, upper panel by applying a growth curve to the data.

#### Gene groups

We divided the proteome into 12 groups, 11 of which were based on SGD GO annotations or KEG annotations, which together account for 80% of the proteome (by protein abundance). The rest were grouped together as an additional 12^th^ group. There is a small overlap between the groups, See Supplementary Table 1 for the protein names in each group.

#### RNAseq transcription protocol and analysis

As described in (Voichek et al., 2016). There are 6-8 repeats of each strain in each condition.

#### Published RNA datasets

The different carbon data sets (Figure 2B, Figure 2- -figure supplement 1) were obtained from (Gasch et al., 2000). Data for strains growing on xylulose (Figure 2B, upper panel) was obtained from (Tamari et al., 2014, 2016). We compared ribosomal fractions from the RNAseq data with ChIP data between the two papers, and excluded from the analysis strains with high variability. Deletion strains (Figure 2B, bottom panel) datasets were obtained from (Kemmeren et al., 2014; O’Duibhir et al., 2014).

#### RNA data of cells growing in Low Pi media (0.06mM)

Logarithmic wild-type cells (≈0.4 OD) grown in SC medium were washed in low Pi (0.06mM) media. Cells were then diluted and inoculated to fresh low Pi media to an initial OD of 0.05. Cells were grown in 30°C for several hours. In every time point, as indicated at Figure 4-figure supplement 1A, a sample was extracted and RNAseq was performed as described above.

#### Ribosomal fraction analysis in the RNA data

The ribosomal fraction was calculated by dividing the number of mRNA reads of ribosomal proteins by the total reads. In the ChIP data, the ribosomal fraction was calculated as the relative amount normalized to the SC condition (Logarithmic mean).

#### mCherry levels measurements

The mCherry levels were calculated from our mass-spectrometry data. To compare the mCherry between the different copy numbers, we used the relative mass-spectrometry data. In order to measure our reference strain (one copy in SC), we used TDH3p-mCherry (protein fusion) strain as described in the Strains section. The amount of mCherry in this strain is equal to the TDH3 protein (≈2.2% base on the iBAQ parameter). The mCherry level of our one mCherry copy strain was 14.5% lower than the mCherry fluorescence of the TDH3p-mCherry strain. Thus, the mCherry level of one copy strain is ≈1.9 % of the proteome. Notably, this calibration is in very good agreement with our previous fluorescence based calibration (Kafri et al., 2016b).

#### Live microscopy environment perturbation experiment

Cells were grown in a standard FCS2 flow cell (Bioptechs) improved similar to (Charvin et al., 2010). Cells growing in the flow cell were imaged with Olympus LX81 microscope with automated stage and ZDC autofocus and cooled CCD camera (Hamamatsu). In order to grow the cells in a planar layer while simultaneously control their extra cellular environment, standard FCS2 flow cell (Bioptechs) was used and improved in the following way (Charvin et al., 2010): the 40 mm round cover slips were coated with a thin layer of PDMS (Sylgrad 184, GE). This was done in a clean room, using a suitable spinning procedure to achieve layers of ≈30 μm thick. To confine the cells and prevent their movement, while medium is flowing through the chamber (Balaban et al., 2004), a diffusive cellulose membrane, which was cut from dialysis tubes (Sigma Aldrich, D9527) was used.

The membrane was cleaned and prepared as described in (Charvin et al., 2010). The cells are confined between the PDMS layer and the membrane, while the medium is flowing above the membrane. In this way the medium reaches the cells area without exerting too much force on the cells. The flow cell is then connected to two micro-perfusion pumps (Instech), which are controlled automatically by the computer via a D/A convener (measurement computing usb-3110), enabling the exchange of medium during the experiment without perturbing the cells. The media tanks for the environment perturbation experiment were continuously stirred, to allow for better aeration of the medium. The whole system is controlled by ImagePro 6.3.1 (Media Cybernetics).

Galactose to glucose shift experiments were done by replacing synthetic complete medium based on 2% galactose (SC-Gal) with synthetic complete medium based on 2% glucose (SC-Glu). Cell volume was estimated from the bright field images assuming that the yeast cells are prolate spheroids.

#### Polysome Profiling

100ml cultures of exponentially growing yeast cells (OD_600_ 0.2-0.4 with at least 6 exponential divisions since their dilution) were treated with 100μg/ml Cycloheximide for 5 minutes at 30°C, harvested, and then washed with cold Buffer AB (20mM Tris-HCL pH 7.4, 50mM KCL, 10mM MgCL2, 1 mM DTT and 100μg/ml Cycloheximide). The cells were lysed with 250 ml Buffer AB using glass beads as in (Pospísek and Valásek, 2013). Cleared lysates were loaded onto 10-50% sucrose gradients with the same AB buffer composition and containing 100μg/ml Cycloheximide, and centrifuged at 39,000 RPM in a SW41 rotor for 2.5 hours at 4°C. Gradients were continuously recorded using Biocomp gradient station at OD_254_ nm.

Analysis of gradients was carried out in Matlab with a customized script. Briefly, profiles were aligned in the x-axis by their peak locations and in the y-axis to SC low burden. A blank sucrose gradient measurement was subtracted from all the samples. Next, the integrals of the 40S, 60S, monosomes and polysome fractions (disomes and higher) were measured. The ratio of polysomes to monosomes was defined as the (polysomes integral) / (40S+60S+monosomes integrals).

